# Prevailing homozygous deletion of interferon and defensin genes in human cancers

**DOI:** 10.1101/101741

**Authors:** Zhenqing Ye, Haidong Dong, Ying Li, Haojie Huang, Han Liang, Jean-Pierre A. Kocher, Liguo Wang

## Abstract

Interferons and defensins are antimicrobial peptides that can also induce anti-tumor immunity. By analyzing the copy number profiles of 10,759 patients across 31 cancer types, we found the homozygous deletions of interferon and defensin genes are prevailing in most human cancers, and that patients with these homozygous deletions exhibited significant reduced overall survival or disease-free survival. We further demonstrated that the homozygous deletion of interferon and defensin genes significantly impacted the expression of genes regulated by tumor necrosis factor (TNF) and IFNγ. Our findings suggested a novel immune escape mechanism that disrupts the tumor cells’ ability to be recognized, and have implications for personalized immunotherapy.

## Abbreviation of TCGA cancer types

ACC: Adrenocortical Carcinoma
BLCA: Bladder Urothelial Carcinoma
BRCA: Breast Invasive Carcinoma
CESC: Cervical Squamous Cell Carcinoma and Endocervical Adenocarcinoma
CHOL: Cholangiocarcinoma
COAD: Colon Adenocarcinoma
DLBC: Diffuse Large B-cell Lymphoma
ESCA: Esophageal Carcinoma
GBM: Glioblastoma Multiforme
HNSC: Head and Neck Squamous Cell Carcinoma
KICH: Kidney Chromophobe
KIRC: Kidney Renal Clear Cell Carcinoma
KIRP: Kidney Renal Papillary Cell Carcinoma
LAML: Acute Myeloid Leukemia
LGG: Brain Lower Grade Glioma
LIHC: Liver Hepatocellular Carcinoma
LUAD: Lung Adenocarcinoma
LUSC: Lung Squamous Cell Carcinoma
MESO: Mesothelioma
OV: Ovarian Serous Cystadenocarcinoma
PAAD: Pancreatic Adenocarcinoma
PCPG: Pheochromocytoma and Paraganglioma
PRAD: Prostate Adenocarcinoma
READ: Rectum Adenocarcinoma
SARC: Sarcoma
SKCM: Skin Cutaneous Melanoma
STAD: Stomach Adenocarcinoma
TGCT: Testicular Germ Cell Tumors
THCA: Thyroid Carcinoma
THYM: Thymoma
UCEC: Uterine Corpus Endometrial Carcinoma
UCS: Uterine Carcinosarcoma
UVM: Uveal Melanoma

## Introduction

Somatic copy number alteration (SCNA) is one of the major sources of genome instability, which plays critical roles in tumorigenesis. Previous studies estiamte that 25% of the cancer genome is affected by arm-level SCNAs, and 10% by focal SCNAs with 2% overlap^1^. Mapping the focal SCNAs that are recurrent in different tumor types could potentially reveal the molecular mechanisms of tumorigenesis and identify therapeutic targets. Through the analysis of the copy number profiles of 4934 cancers across 11 cancer types, The Cancer Genome Atlas (TCGA) project identified 140 recurrent focal SCNAs including 70 amplified regions (959 affected genes) and 70 deleted regions (2084 affected genes)^2^. Another study explored the copy number profiles of 3131 cancers across 26 cancer types and identified 158 recurrent focal SCNAs including 76 amplification (1566 affected genes) and 82 deletions (2001 affected genes). These large-scale SCNA profiling studies significantly improved our understandings of the genomic landscapes of human tumors, and have identified hundreds of oncogenes and tumor suppressors located in amplified and deleted regions, respectively. However, It is still a challenge to pinpoint the common biological pathways given the large number of genes affected by the focal SCNAs.

The loss of tumor suppressor genes plays important role in cancer biology. Tumor suppressor genes generally follow the “two-hit hypothesis”, which suggests that both alleles of the same gene must be deactivated before an negative effect is observed^3^. Therefore, homozygously deleted genes provide an important resource for identifying cancer-causing tumor suppressors. Although similar work has been done in cell lines^4^, large-scale and genome-wide analysis have not been conducted in of patient specimens. In this study, we re-analyzed the TCGA copy number profiles of 10,759 patients across 31 cancer types to identify recurrent, homozygously deleted genes, aiming at identifying common biological pathways implicated in tumorigenesis. Strikingly, except for eight well-known tumor suppressors such as *PTEN* and *RB1*, we found that all identified HDGs were located in only two loci–8p21-23 and 9p21. In particular, we found interferon gene cluster (located in 9p21) and defensin gene cluster (located in 8p21-23) were homozygously deleted in at least 7% of patients in 19 out of 31 (61%) cancer types. Survival analyses in different tumor types indicated that patients with homozygous deletion of interferons or defensins exhibit dramatically reduced overall survival or disease-free survival. Since a large body of evidence suggest that interferons and defensins have a major role in tumor immunity by recognizing tumor cells and serve as a bridge to spontaneous adaptive T cell response^5-11^, our findings suggested a common molecular mechanisms mediated by the loss of interferons and defensins, through which tumor cells escape immune detection, and provided solid evidence supporting the “evading immune destruction” as the new emerging cancer hallmark^12^.

## Results

To identify homozygously deleted genes that are common to human cancers, we analyzed the CNV profiles of 10,759 patients across 31 cancer types generated from TCGA Research Network (**Table S1**). We first calculated the homozygous deletion frequency for each gene in each cancer type (**Table S2**). Then, we identified 242 genes whose average homozygous deletion frequency across 31 cancer types is larger than 3%. Strikingly, we found 234/242 genes (96.7%) were located on 8p21-23 or 9p21, with only 8 genes (3.3%) located in other regions (**Table S3**). As expected, 7 out of these 8 genes are known tumor suppressors including *PTEN*, *RB1*, *DMD*, *PTPRD*, *PDE4D*, *WWOX* and *LRP1B*, and all these 7 genes were identified as the potential targets of focal SCNAs by two independent studies^1,2^. The molecular function of another gene *CCSER1* (alias *FAM190A*) is largely unknown, but its homozygous deletion has been frequently observed in human cancers^13-15^. Functional classification of the 234 genes revealed that interferons (16 genes, q-value = 8.37×10^-20^) and defensins (24 genes, q-value = 1.65×10^-30^) are the most significantly enriched terms (**Table S4**). The 16 type I interferon genes are located on 9p21, which includes 13 IFN-α genes, 1 IFN-β, 1 IFN-ε and 1 IFN-ω gene. The 24 defensin genes are located on 8p21-23, which includes 6 α-defensin and 18 β-defensin genes. Since both interferons and defensins are involved in innate immune response and play important roles in recognizing tumor cells and inducing anti-tumor immune response, the recurrent, homozygous deletion of these genes suggested a common molecular mechanism through which tumor cells escape immune destruction.

We observed the homozygous deletion of interferons (HDIs) and defensins (HDDs) from all 31 cancer types except PCPG (pheochromocytoma and paraganglioma), THCA (thyroid carcinoma) and LAML (acute myeloid leukemia) (**Fig. 1**). Using a 5% alternation frequency as a threshold, we defined 12/31 (38.7%) tumors as the low HDI/HDD group (or L-Type tumors) and 19/31 (61.3%) tumors as the high HDI/HDD group (**Fig. 1**, **Table S5**). Interestingly, we found that 50% (6/12) of low HDI/HDD tumors were rare cancer types according to TCGA’s classification (https://cancergenome.nih.gov/cancersselected/RareTumorCharacterizationProjects), but only 15.8% (3/19) of high HDI/HDD tumors were rare cancer types (*P* = 0.056, two-sided Fisher’s exact test) (**Fig. 1**, **Table S5**). This contrast pattern suggested that HDI/HDD might be a common molecular mechanism contributing to the prevalence of major cancer types. In the high HDI/HDD group, two brain tumor types LGG (brain lower grade glioma) and GBM (glioblastoma multiforme) exhibited the lowest (7.2%) and highest (30.5%) alternation frequencies, respectively. We further divided the high HDI/HDD group into three subtypes using the prevalence ratio (*PR* = {# of HDIs}/{# of HDDs}) of 5 as the threshold (**Fig. 1**, **Table S5)**: The I- type was defined as cancer types with HDIs at least 5 times more prevalent than that of HDDs (*PR* ≥ 5), including GBM, LGG, MESO, and PAAD (**Fig. 2A, Fig. S1-2**). The D type was defined as cancer types with HDDs at least 5 times more prevalent than that of HDIs (*PR* ≤ 0.2), including COAD, READ, LIHC, UCS and PRAD (**Fig. 2B, Fig. S3-4**). The C type referred to those cancer types with both HDIs and HDDs (0.2 < *PR* < 5), including BRCA, BLCA, ESC, HNSC, LUAD, LUSC, OV, SARC, SKCM and STAD (**Fig. 2C, Fig. S5-8**).

**Figure 1.**
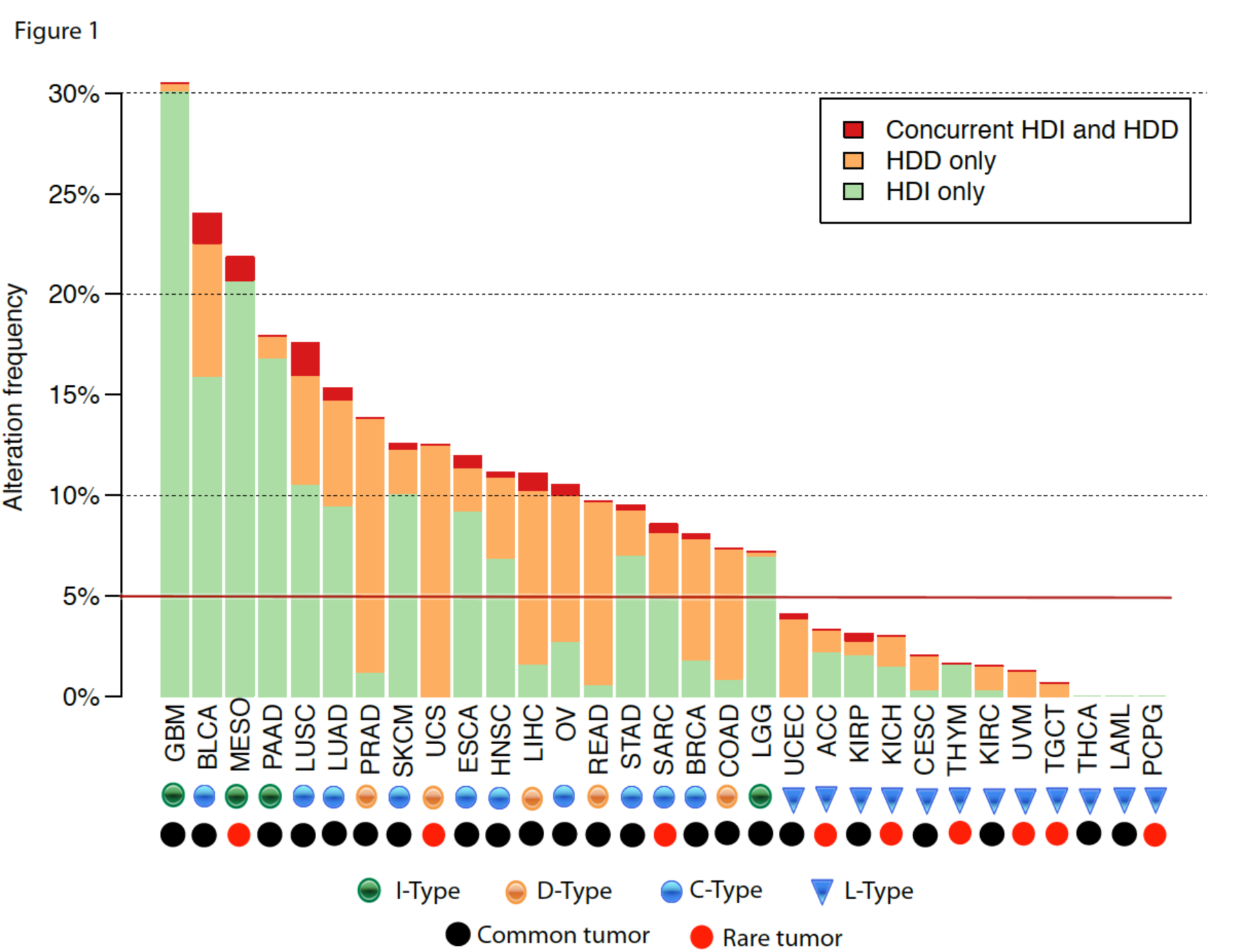
Homozygous deletion frequencies of interferon and defensin genes in 31 cancer types. Within each cancer type, patients were divided into three groups: homozygous deletion of interferon genes (HDI) only (green), homozygous deletion of defensin genes (HDD) only (orange) and concurrent HDI and HDD (red). The 31 cancers were classified into 4 types: L-type refers to cancer types with overall homozygous deletion frequency of HDI and HDD less than 5% (blue triangle); I-type refers to cancer types in which HDI is dominant (green circle); D-type refers to cancer types in which HDD is dominant (orange circle); C-type refers to cancer types in which both HDI and HDD are prevalent (blue circle). Common and rare tumors are designed by TCGA and indicated using black and red circles, respectively.

**Figure 2.**
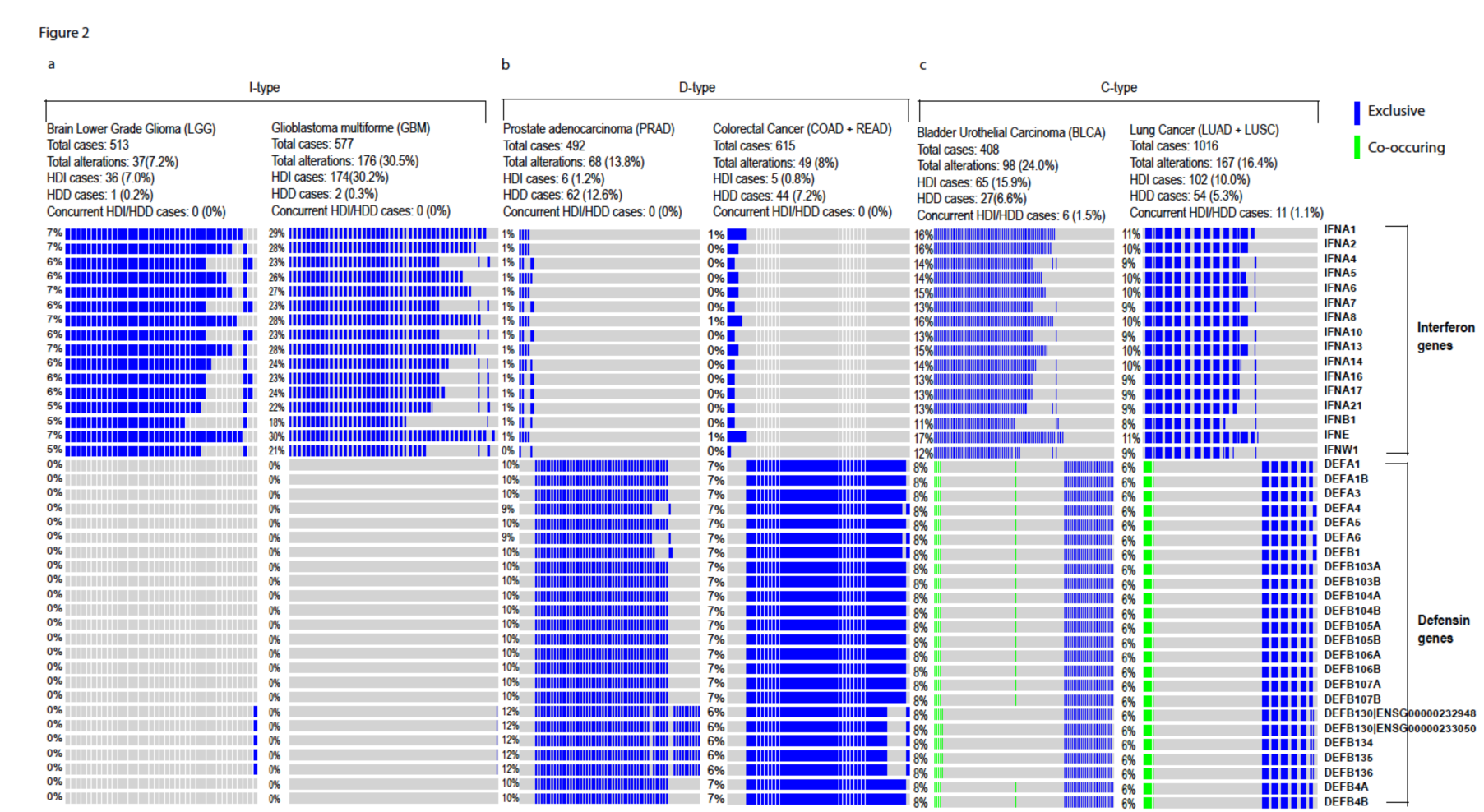
Oncoprint plots showing gene level alteration patterns of interferon and defensin genes. Each row represents a gene and each column represents a patient. Blue vertical bars indicate patients with either HDD or HDI, green vertical bars indicate patients with concurrent HDD and HDI. (A) I-type cancers using Brain Lower Grade Glioma and Glioblastoma multiforme as examples. (B) D-type cancers using prostate adenocarcinoma and colorectal cancer as examples. (C) C-type cancers using bladder urothelial carcinoma and lung cancer as examples. Colorectal cancer is the combined cohort of colon adenocarcinoma and rectum adenocarcinoma. Lung cancer is the combined cohort of lung adenocarcinoma and lung squamous cell carcinoma.

Since both interferons and defensins play critical roles in anti-tumor immunity, we next investigated if patients whose genome contains HDI or HDD lesions exhibited worse overall survival (OS) or disease-free survival (DFS). Indeed, HDI or HDD patients showed significantly reduced OS or DFS in multiple cancer types (**Fig. 3**). For examples, compared to LGG patients without HDI or HDD (median overall survival time 93.13 months), the median overall survival time of patients carrying HDI or HDD was reduced by 74% to 24.38 months (LogRank test *P*-value = 0) (**Fig. 3A**). Similarly, the median disease free survival time for bladder cancer patients was reduced by 60% from 43.96 months to 17.51 months (LogRank test *P*-value = 0.0026) (**Fig. 3E**), and the median overall survival time of lung cancer patients was reduced by 30% from 54.30 months to 37.91 months (LogRank test *P*-value = 0.0085) (**Fig. 3F**). The striking associations of HDI/HDD status with worse clinical outcomes across cancer types suggested their remarkable prognostic values.

**Figure 3.**
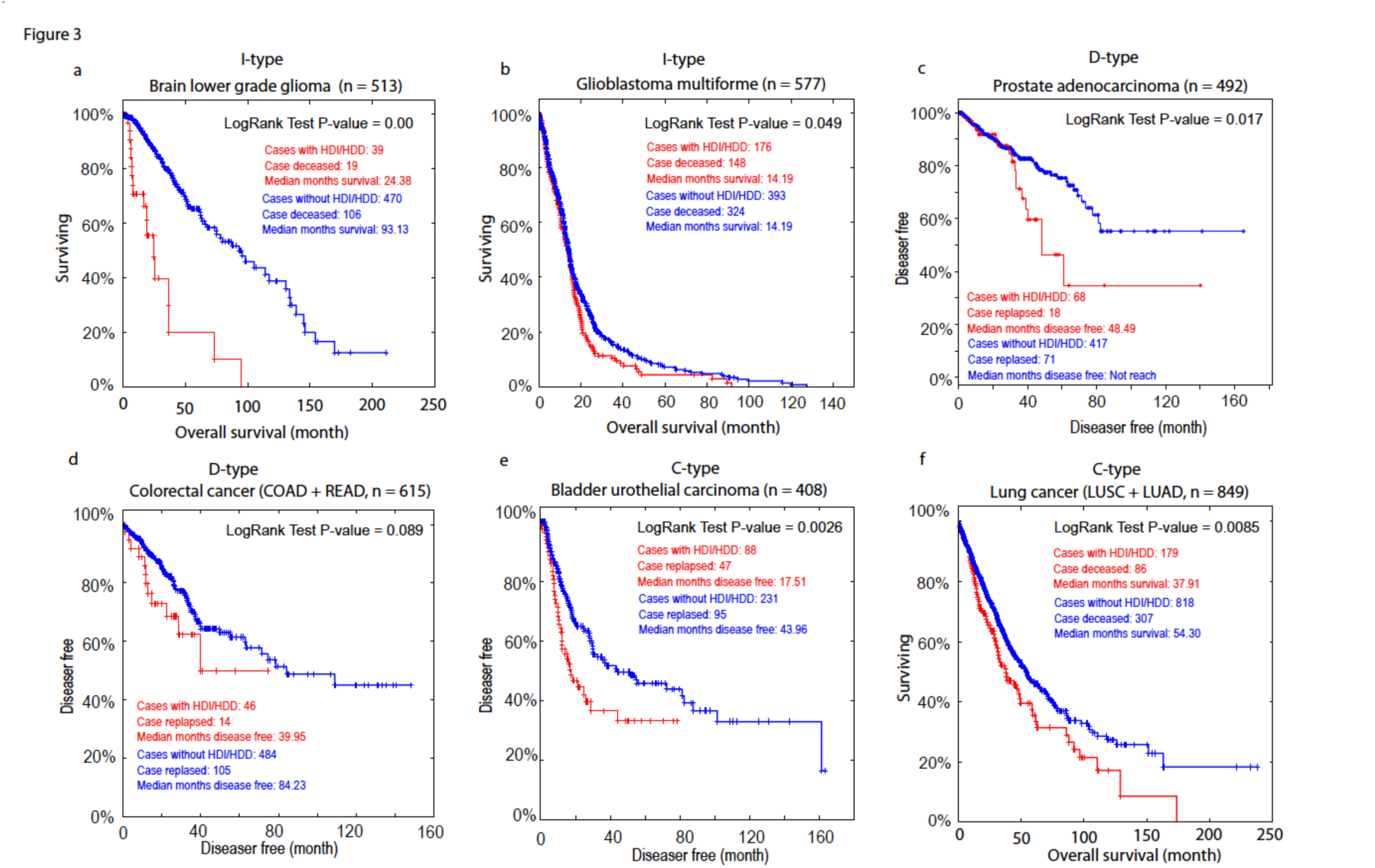
Comparing overall survival or disease-free survival between patients with (red) and without (blue) HDIs/HDDs. Survival curves for Brain Lower Grade Glioma, Glioblastoma multiforme, Prostate adenocarcinoma, Colorectal cancer, Bladder Urothelial Carcinoma and Lung cancer are indicated in panel (A) - (F), respectively. Colorectal cancer is the combined cohort of colon adenocarcinoma and rectum adenocarcinoma. Lung cancer is the combined cohort of lung adenocarcinoma and lung squamous cell carcinoma.

We found that interferon genes are located approximately 485 Kb to 890 Kb away from *CDKN2A* (Cyclin Dependent Kinase Inhibitor 2A) (**Table S3**), a well-known tumor suppressor. The homozygous deletion of these genes have been reported in mesothelioma and numerous cancer cell lines^4,16,17^. Similarly, defensin genes are located ~5Mb away from another tumor suppressor *CSMD1* (CUB And Sushi Multiple Domains 1)^18^ (**Table S3**). We asked the question whether homozygous deletions of interferon and defensin gene clusters were passive events hitchhiked by their nearby known tumor suppressors, or they played an active role in tumorigenesis and affected the patient survival. Therefore, we tested if the loss of interferon and defensin genes could significantly impact gene expression. We performed gene expression analysis on each of the 17 cancer types (COAD and READ were combined as colorectal cancer, LUSC and LUAD were combined as lung cancer) (**Fig. 1**, **Table S5**). Through differential gene expression analysis of patients with and without HDI/HDD lesions in each tumor type, we detected 4599 genes whose expression were significantly (FDR < 0.01) altered in at least one cancer type (**Table S6**). For example, *KLHL9*, a gene located within the interferon gene cluster, was identified as significantly down-regulated in 12 out of 17 cancer types (**Table S6**). To identify the implicated pathway, we performed Gene Set Enrichment Analysis (GSEA) for the top 143 genes whose expressions were significantly changed in at least 5 cancer types (**Table S6, Fig. S9**)^19^. GSEA results showed that the “immune system” was the only gene set that enriched in these 143 genes (FWER corrected *P* = 0.071) (**Fig. S10**). We further performed Ingenuity Pathway Analysis (IPA^®^) analyses for genes differentially expressed between the two groups for each tumor type. Strikingly, we found tumor necrosis factor (TNF) was detected as the top upstream regulator for all the 17 cancer types with extremely significant P-values (**Table 1**). IFN-γ (IFNG) was another common upstream regulator detected in eight tumor types including MESO, PAAD, LGG, Colorectal (COAD + READ), LUNG (LUSC + LUAD), BLCA, SARC and STAD (**Table S6**). Both TNF and IFN-γ have direct and indirect antitumor functions. They are defined as effector molecules of CD4^+^ helper T (Th1) cells, NK cells and CD8^+^ cytotoxic T cells. TNF is also an effector molecule produced from activated macrophages (M1) capable of killing tumors. On the other hand, IFN-γ induces the up-regulation of PD-L1, a immune checkpoint molecule in many human cancer cells. The fact that HDD and HDI specifically impacted the expression of genes involved in immune system demonstrated their active role in tumorigenesis and survival.

**Table 1:**
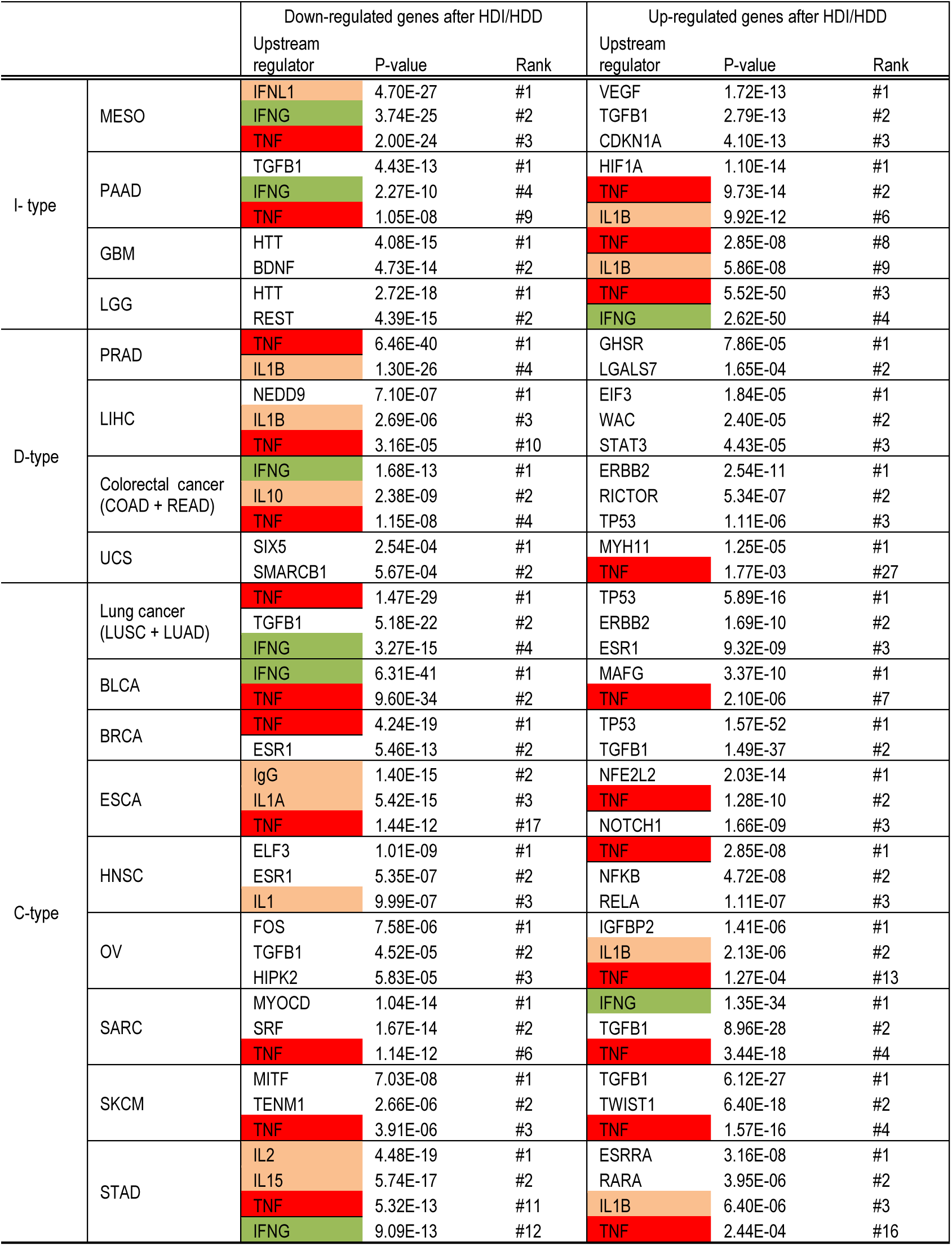
Ingenuity IPA identified top "upstream regulator genes or proteins" from genes that are differentially expressed (FDR < 0.01) in each cancer type.

|  |  | Down-regulated genes after HDI/HDD |  |  | Up-regulated genes after HDI/HDD |  |  |
| --- | --- | --- | --- | --- | --- | --- | --- |
|  |  | Upstream regulator | P-value | Rank | Upstream regulator | P-value | Rank |
| I-type | MESO | IFNL1 | 4.70E-27 | #1 | VEGF | 1.72E-13 | #1 |
|  |  | IFNG | 3.74E-25 | #2 | TGFB1 | 2.79E-13 | #2 |
|  |  | TNF | 2.00E-24 | #3 | CDKN1A | 4.10E-13 | #3 |
|  | PAAD | TGFB1 | 4.43E-13 | #1 | HIF1A | 1.10E-14 | #1 |
|  |  | IFNG | 2.27E-10 | #4 | TNF | 9.73E-14 | #2 |
|  |  | TNF | 1.05E-08 | #9 | IL1B | 9.92E-12 | #6 |
|  | GBM | HTT | 4.08E-15 | #1 | TNF | 2.85E-08 | #8 |
|  |  | BDNF | 4.73E-14 | #2 | IL1B | 5.86E-08 | #9 |
|  | LGG | HTT | 2.72E-18 | #1 | TNF | 5.52E-50 | #3 |
|  |  | REST | 4.39E-15 | #2 | IFNG | 2.62E-50 | #4 |
| D-type | PRAD | TNF | 6.46E-40 | #1 | GHSR | 7.86E-05 | #1 |
|  |  | IL1B | 1.30E-26 | #4 | LGALS7 | 1.65E-04 | #2 |
|  | LIHC | NEDD9 | 7.10E-07 | #1 | EIF3 | 1.84E-05 | #1 |
|  |  | IL1B | 2.69E-06 | #3 | WAC | 2.40E-05 | #2 |
|  |  | TNF | 3.16E-05 | #10 | STAT3 | 4.43E-05 | #3 |
|  | Colorectal cancer (COAD + READ) | IFNG | 1.68E-13 | #1 | ERBB2 | 2.54E-11 | #1 |
|  |  | IL10 | 2.38E-09 | #2 | RICTOR | 5.34E-07 | #2 |
|  |  | TNF | 1.15E-08 | #4 | TP53 | 1.11E-06 | #3 |
|  | UCS | SIX5 | 2.54E-04 | #1 | MYH11 | 1.25E-05 | #1 |
|  |  | SMARCB1 | 5.67E-04 | #2 | TNF | 1.77E-03 | #27 |
| C-type | Lung cancer (LUSC + LUAD) | TNF | 1.47E-29 | #1 | TP53 | 5.89E-16 | #1 |
|  |  | TGFB1 | 5.18E-22 | #2 | ERBB2 | 1.69E-10 | #2 |
|  |  | IFNG | 3.27E-15 | #4 | ESR1 | 9.32E-09 | #3 |
|  | BLCA | IFNG | 6.31E-41 | #1 | MAFG | 3.37E-10 | #1 |
|  |  | TNF | 9.60E-34 | #2 | TNF | 2.10E-06 | #7 |
|  | BRCA | TNF | 4.24E-19 | #1 | TP53 | 1.57E-52 | #1 |
|  |  | ESR1 | 5.46E-13 | #2 | TGFB1 | 1.49E-37 | #2 |
|  | ESCA | IgG | 1.40E-15 | #2 | NFE2L2 | 2.03E-14 | #1 |
|  |  | IL1A | 5.42E-15 | #3 | TNF | 1.28E-10 | #2 |
|  |  | TNF | 1.44E-12 | #17 | NOTCH1 | 1.66E-09 | #3 |
|  | HNSC | ELF3 | 1.01E-09 | #1 | TNF | 2.85E-08 | #1 |
|  |  | ESR1 | 5.35E-07 | #2 | NFKB | 4.72E-08 | #2 |
|  |  | IL1 | 9.99E-07 | #3 | RELA | 1.11E-07 | #3 |
|  | OV | FOS | 7.58E-06 | #1 | IGFBP2 | 1.41E-06 | #1 |
|  |  | TGFB1 | 4.52E-05 | #2 | IL1B | 2.13E-06 | #2 |
|  |  | HIPK2 | 5.83E-05 | #3 | TNF | 1.27E-04 | #13 |
|  | SARC | MYOCD | 1.04E-14 | #1 | IFNG | 1.35E-34 | #1 |
|  |  | SRF | 1.67E-14 | #2 | TGFB1 | 8.96E-28 | #2 |
|  |  | TNF | 1.14E-12 | #6 | TNF | 3.44E-18 | #4 |
|  | SKCM | MITF | 7.03E-08 | #1 | TGFB1 | 6.12E-27 | #1 |
|  |  | TENM1 | 2.66E-06 | #2 | TWIST1 | 6.40E-18 | #2 |
|  |  | TNF | 3.91E-06 | #3 | TNF | 1.57E-16 | #4 |
|  | STAD | IL2 | 4.48E-19 | #1 | ESRRA | 3.16E-08 | #1 |
|  |  | IL15 | 5.74E-17 | #2 | RARA | 3.95E-06 | #2 |
|  |  | TNF | 5.32E-13 | #11 | IL1B | 6.40E-06 | #3 |
|  |  | IFNG | 9.09E-13 | #12 | TNF | 2.44E-04 | #16 |

We further tested if the additional loss of interferon genes could impact patients’ overall survival time, comparing to patients with *CDKN2A* deletion only. For each tumor type, we performed the survival analysis on three patient groups: (I) patients only have *CDKN2A* loss; (II) patients have *CDKN2A* loss and additional interferon genes deletions; and (III) patients have neither *CDKN2A* nor interferon genes deletions. As expected, the overall survival rates of group (II) patients were dramatically reduced as compared to those of group (I) in a variety of tumor types including LGG, GBM, BRCA, lung (LUAD + LUSC), colorectal (COAD + READ), SARC, OV, BLCA and SKCM (**Fig. 4**). The P-values of several tumor types did not reach statistical significance more likely due to limited sample size, for example, only 5 and 14 patients have both *CDKN2A* and interferon genes deletions for colorectal cancer and SARC, respectively. It could be also due to other confounding factors such as PTEN and RB1 deletion, which tend to be mutually exclusive with interferon gene deletions (**Fig. S11**).

**Figure 4.**
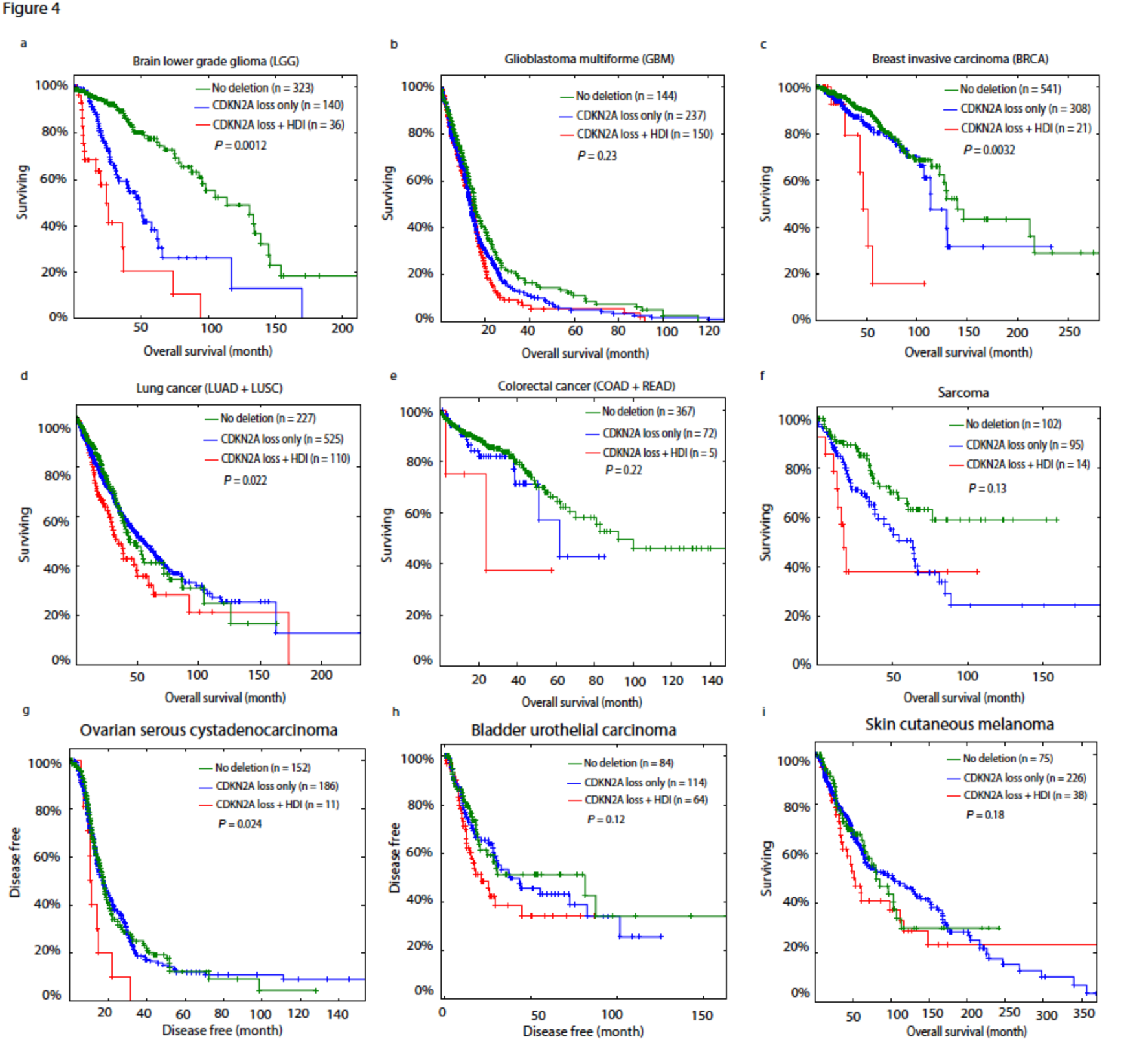
Comparing overall survival between patients have neither CDKN2A nor interferon gene deletions (green), patients only have CDKN2A deletion (blue), and patients have both CDKN2A and interferon gene deletions (red). Survival curves for brain lower grade glioma, glioblastoma multiforme, breast invasive carcinoma, lung cancer, colorectal cancer, sarcoma, ovarian serous cystadenocarcinoma, bladder urothelial carcinoma, skin cutaneous melanoma are indicated in panel (A) – (I), respectively. Colorectal cancer is the combined cohort of colon adenocarcinoma and rectum adenocarcinoma. Lung cancer is the combined cohort of lung adenocarcinoma and lung squamous cell carcinoma. Log rank test P-values were calculated by comparing patients only have CDKN2A deletion (blue), and patients have both CDKN2A and interferon gene deletions (red).

## Discussion

Evading immune destruction is considered as one of the two new emerging hallmarks of cancer^12^. A growing body of evidence has suggested that interferons and defensins play important roles in recognizing tumor cells and inducing immune responses. For example, type I interferons are critical for the innate immune to recognize a growing tumor and activate CD8+ T cell responses^9^, and that intratumoral production of type I interferon after innate immune recognition of tumor cells is critical for activating natural adaptive immune response against tumors *in vivo*^20^. Defensins can promote anti-tumor adaptive immune responses in mice^21^, *in vitro* tumor cell cytolysis^7^, and induce antitumor immunity when fused with nonimmunogenic tumor antigen and enhance antibody responses^22,23^. In this study, we analyzed the SCNA profiles of 10,759 cancer tissues and found homozygous deletions of interferon and defensin gene clusters were highly recurrent in 19 cancer types with alternation frequency ranging from 7.2% to 30.5%, which is generally much higher than that of PTEN and RB1 in the same cancer type (**Fig. S11**). More importantly, patients with HDI or HDD lesions exhibited significantly reduced overall or disease-free survival in a variety of tumors. Given that both interferons and defensins play critical roles in initiating tumor immunity, the high prevalence of HDIs and HDDs in human cancers indicated that this might be the generic mechanism through which tumor cells avoid anti-tumor destruction after tumorigenesis.

Immunotherapies targeting PD-1:PD-L-1 interaction have been demonstrated to be efficacious in a number of cancer types^24–26^, however, therapeutic resistance is frequently observed and the mechanisms of both *de novo* and acquired immune-resistance are mostly unknown. Defects in the interferon signaling pathway have been proposed as a potential mechanism of cancer escape (insensitivity) to immunotherapy in mice and prostate cancer cell line^27,28^. In mice, type I interferon signals are required to initiate the antitumor CD8^+^ response, and mice without INF-α/β receptor cannot reject immunogenic tumor cells^9,10^. Consistent with these preclinical observations, our observation that type I interferon genes were frequently deleted in 14 tumor types (I-type and C-type) suggested a generic mechanism through which tumors develop acquired immuno-resistance, and revealed new ‘omics-’ based biomarkers to identify responsive patients. Our results also suggest that personalized immunotherapies based on patient’s interferon and/or defensin deletion status could be considered.

When comparing gene expression profiles of patients having HDI/HDD with those without HDI/HDD events, all the 16 interferon and the 24 defensin genes were not identified as differentially expressed except for *DEFB1* which is significantly down regulated in ESCA, LIHC and OV (**Table S6**). The major reason is that most interferon and defensin genes are not normally active in these tissues but can be activated during oncogenic process (**Fig. S12**). For example, although DEFA1-3 are primarily expressed in neutrophils and NK cells, up-regulation of HNP1-3 (encoded by DEFA1-3, respectively) have been detected in tumor tissues in colorectal and other cancers ^8,29–31^. Up-regulation of HNP1-3 in tumor tissues might originate from tumor infiltrating immune cells, however, several studies performed on cell lines demonstrated that tumor cells could produce HNP1-3 by themselves^8,32^. Similarly, IFN-α and IFN-β genes can be produced from both tumor cells and infiltrating innate immune cells to elicit anti-tumor immune response^33^. Given the role of interferons and defensins in recognizing tumor cells and inducing immune response, we hypothesize that their homozygous deletion could confer to tumor cells a growth/survival advantage after tumorigenesis. Further analysis is warranted to validate this point.

We found that patients of I-type tumors almost exclusively have HDI but not HDD (**Fig. 2A, Fig. S1-2**), and patients of D-type almost exclusively have HDDs but not HDIs (**Fig. 2B, Fig. S3-4**). Even for C-type tumor such as BRCA (n = 1079), there were 20 (1.9%) patients having HDIs and 65 (6.0%) patients having HDDs but only 2 (0.2%) patients with concurrent HDI and HDD (**Fig. S5**). These data suggested HDI and HDD had tendency towards mutual exclusivity both within and across tumors, even though they did not reach statistical significance likely due to limited number of cases. This observation is consistent with the concept that alterations within the same pathway are often mutually exclusive^34,35^. One open question is whether patients with concurrent HDI and HDD exhibit worse or favorable clinical outcomes. The low occurrence of patients with concurrent HDI and HDD in TCGA cohort preclude rigorous statistical analysis. However, analysis of LUSC cohort that has the highest concurrent HDI and HDD frequency (n=8), we found the diagnosis age of patients with concurrent HDI and HDD (median = 60) is much smaller than other patients (median = 68) with Wilcoxon two-sided p-value 0.079 (**Fig. S13**), suggesting concurrent HDI and HDD is associated with early tumor onset in LUSC. However, a much larger cohort is needed to test this hypothesis in LUSC and other tumors.

## Methods

### The Cancer Genome Atlas Copy Number Variation Data and Analysis

Thresholded copy number variation data of 10,843 patients (33 cancer types) were downloaded from the Cancer Genome Hub at the University of California at Santa Cruz (https://genome-cancer.ucsc.edu/)^36^. After removing CHOL (n = 36) and DLBC (n = 48) cohorts that have less then 50 patients, SCNA profiles of 10,759 patients were analyzed. These 10,759 patients composed 31 cancer types, which include 22 common cancers and 9 rare cancers. Common and rare tumor designation is according to http://cancergenome.nih.gov/cancersselected/RareTumorCharacterizationProjects.

Matched overall survival or disease-free survival data were downloaded from CBioPortal (http://www.cbioportal.org/)^37^. In brief, Gistic2 generated gene level CNV estimates were thresholded into discrete values -2, -1, 0, 1, 2 representing homozygous deletion, single copy deletion, diploid neutral, low copy number gain and high copy number amplification, respectively. Homozygous deletion frequency is calculated as:

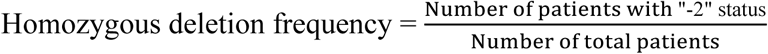

Functional annotation of the homozygously deleted genes was performed by DAVID (https://david.ncifcrf.gov/)^38^ and ConsensusPathDB gene set overrepresentation analysis (http://cpdb.molgen.mpg.de/CPDB)^39^. Both tools produced the similar results. The p-values and q-values in **Table S4** were produced by ConsensusPathDB.

### Gene expression analysis

For each tumor type, cancer patients were first divided into two groups according to the HDI and/or HDD status. Patients without RNA-seq data were removed. Gene level expression estimates (i.e. RSEM normalized read count) were downloaded from TCGA Data Portal (https://tcga-data.nci.nih.gov/). EdgeR was used to compare gene expression profiles between patients with HDI/HDD and those without HDI/HDD. Differentially expressed genes were determined by the p-value cutoff 0.01^40^.

### Gene set enrichment analysis

A total of 143 genes whose expression significantly (p-value < 0.01) changed in at least 5 tumor types were selected for Gene Set Enrichment Analysis (GSEA). These 143 genes were pre-ranked by the average fold change among 17 tumor types before running GseaPreranked module. Used gene sets include Biocarta (v5.2), KEGG (v5.2), Reactome (v5.2). Gene sets larger than 500 or smaller than 15 were removed.

### Survival and statistical analysis

Cancer patients were divided into two groups according to the HDI and/or HDD status. Survival analysis was performed using the “survival” R package available from http://cran.r-project.org. The Log-rank test was used to evaluate if the difference of overall survival or disease-free survival between the above two subgroups was statistically significant. CoMEt exact test was used to evaluate the mutual exclusivity of HDIs and HDDs in each tumor type^41^.

**Author Contributions:** Dr Wang and Ye had full access to the data in this study and take the responsibility for the integrity of the data and the accuracy of the analysis.

Study concept and design: Wang, Kocher, Liang

Survival analysis: Ye Acquisition, analysis and interpretation of the data: Ye, Li and Wang

Critical revision of the manuscript for important intellectual content: Dong, Huang, Liang and Kocher

**Conflict of Interest Disclosures:** None reported

**Funding/Support:** This study was supported by the Center for Individualized Medicine (Pharmacogenomics Program), Mayo Clinic.

## supplementary figures

**Figure S1-S8.**
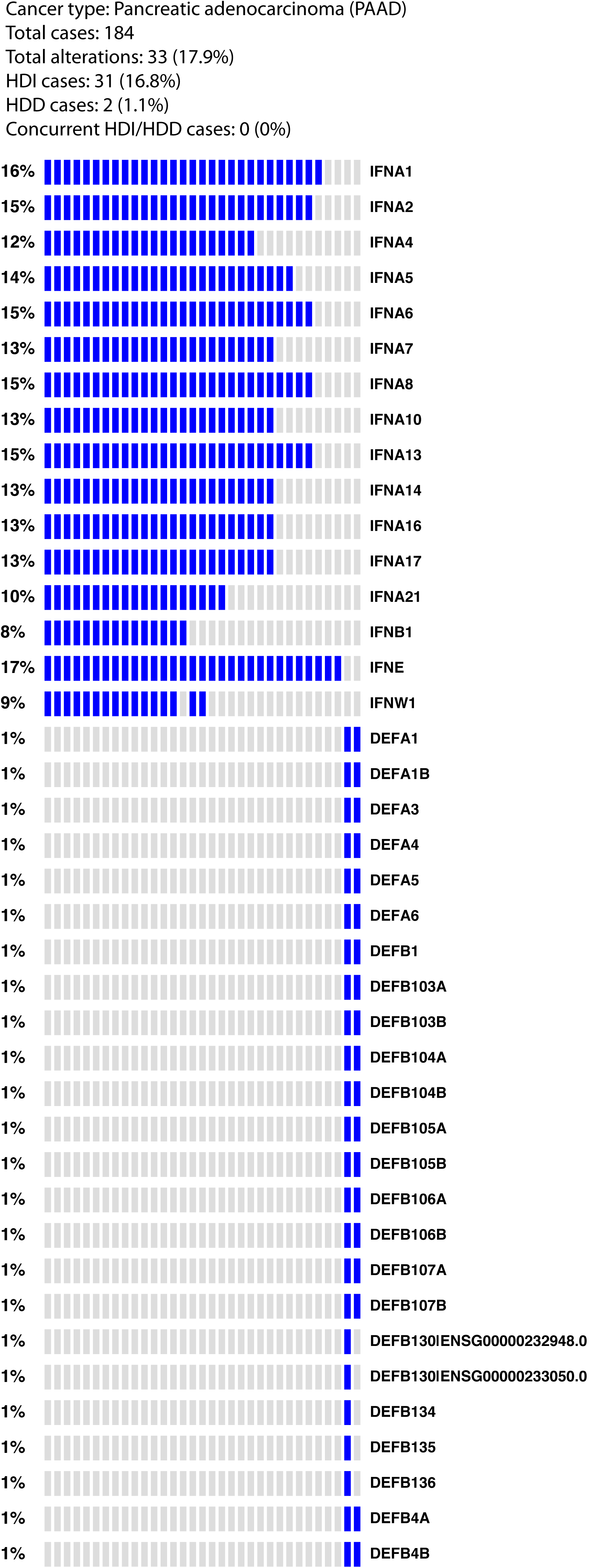
Oncoprint plots for Pancreatic adenocarcinoma (PAAD), Mesothelioma (MESO), Liver hepatocellular carcinoma (LIHC), Uterine carcinosarcoma (UCS), Breast invasive carcinoma (BRCA), Ovarian serous cystadenocarcinoma (OV), Head and Neck squamous cell carcinoma (HNSC), and Skin Cutaneous Melanoma (SKCM).

**Figure S2.**
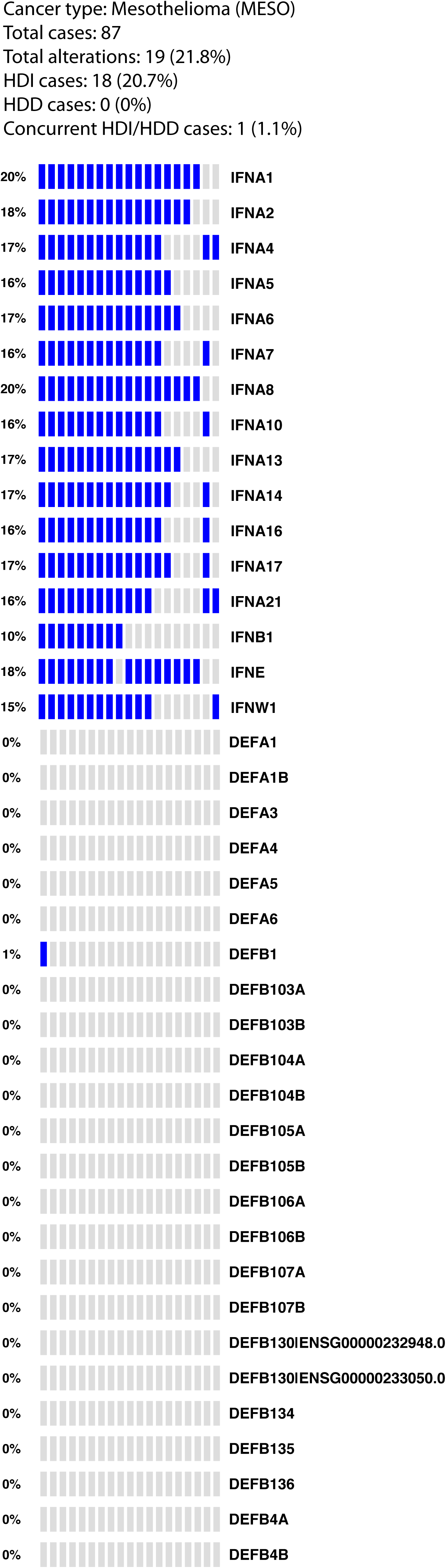
Oncoprint plots for Pancreatic adenocarcinoma (PAAD), Mesothelioma (MESO), Liver hepatocellular carcinoma (LIHC), Uterine carcinosarcoma (UCS), Breast invasive carcinoma (BRCA), Ovarian serous cystadenocarcinoma (OV), Head and Neck squamous cell carcinoma (HNSC), and Skin Cutaneous Melanoma (SKCM).

**Figure S3.**
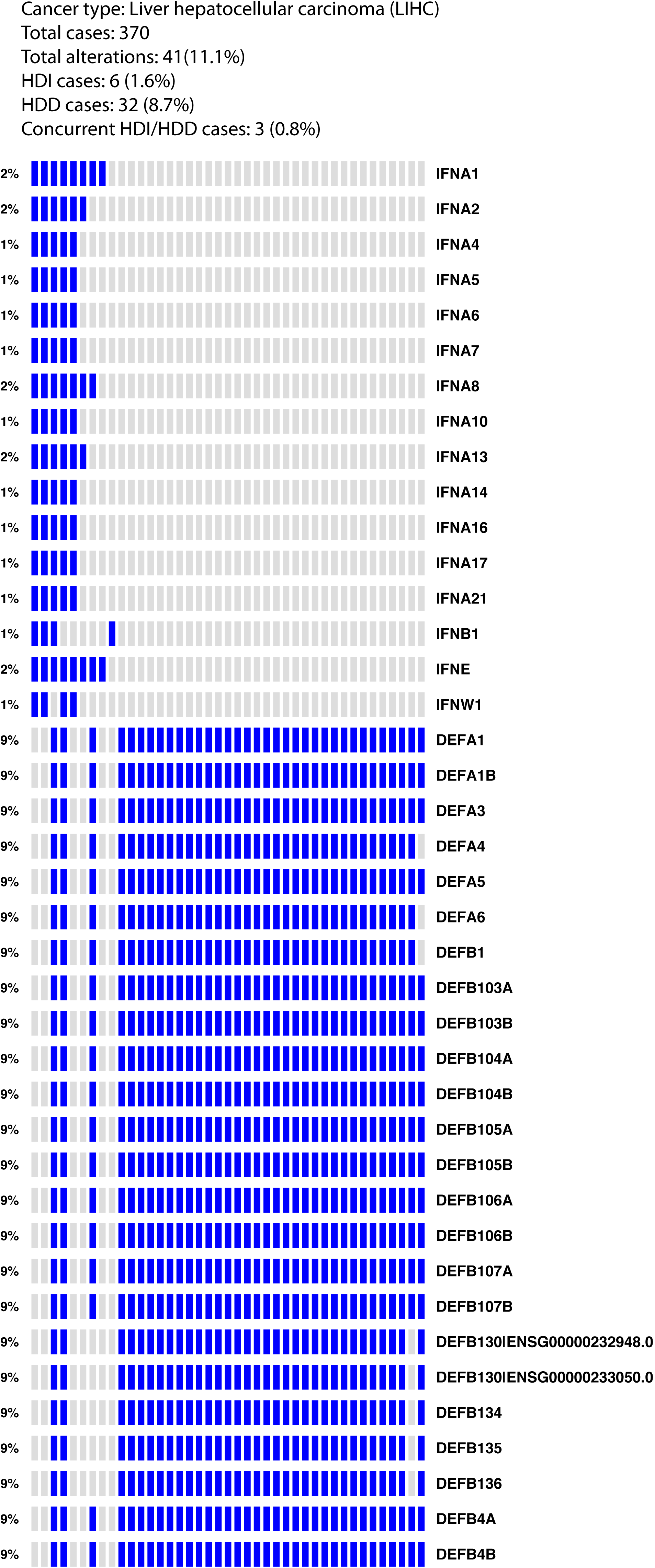
Oncoprint plots for Pancreatic adenocarcinoma (PAAD), Mesothelioma (MESO), Liver hepatocellular carcinoma (LIHC), Uterine carcinosarcoma (UCS), Breast invasive carcinoma (BRCA), Ovarian serous cystadenocarcinoma (OV), Head and Neck squamous cell carcinoma (HNSC), and Skin Cutaneous Melanoma (SKCM).

**Figure S4.**
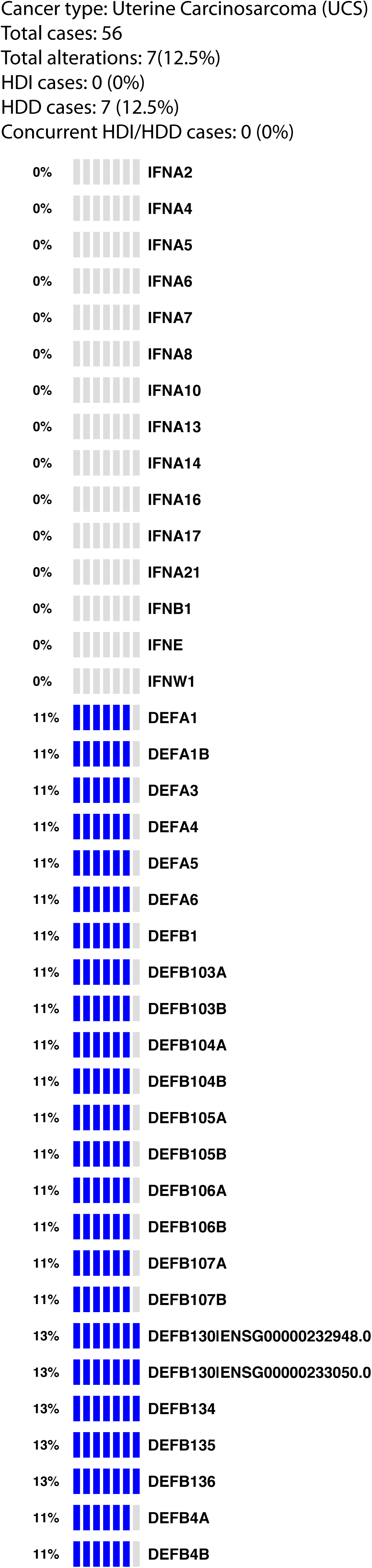
Oncoprint plots for Pancreatic adenocarcinoma (PAAD), Mesothelioma (MESO), Liver hepatocellular carcinoma (LIHC), Uterine carcinosarcoma (UCS), Breast invasive carcinoma (BRCA), Ovarian serous cystadenocarcinoma (OV), Head and Neck squamous cell carcinoma (HNSC), and Skin Cutaneous Melanoma (SKCM).

**Figure S5.**
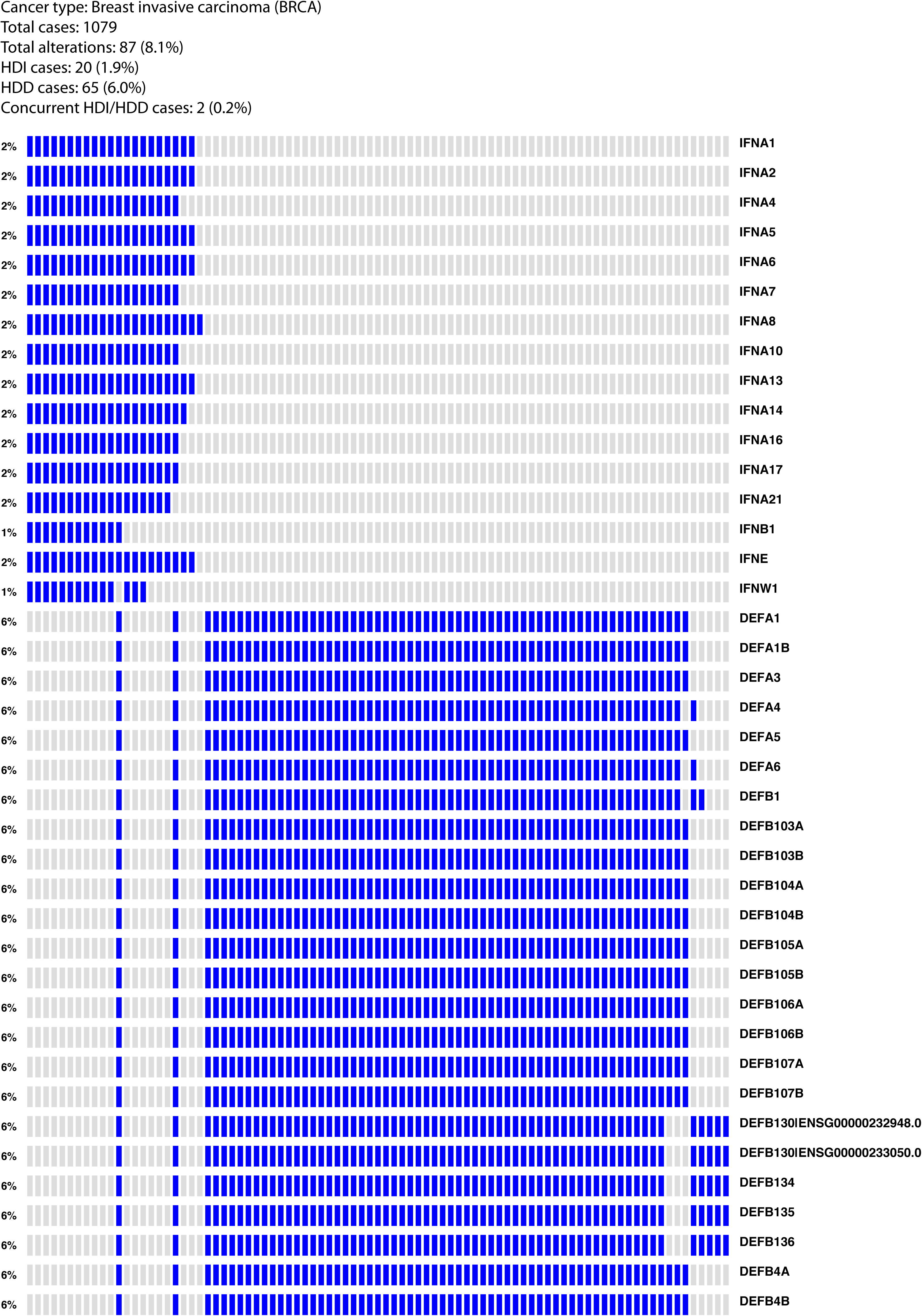
Oncoprint plots for Pancreatic adenocarcinoma (PAAD), Mesothelioma (MESO), Liver hepatocellular carcinoma (LIHC), Uterine carcinosarcoma (UCS), Breast invasive carcinoma (BRCA), Ovarian serous cystadenocarcinoma (OV), Head and Neck squamous cell carcinoma (HNSC), and Skin Cutaneous Melanoma (SKCM).

**Figure S6.**
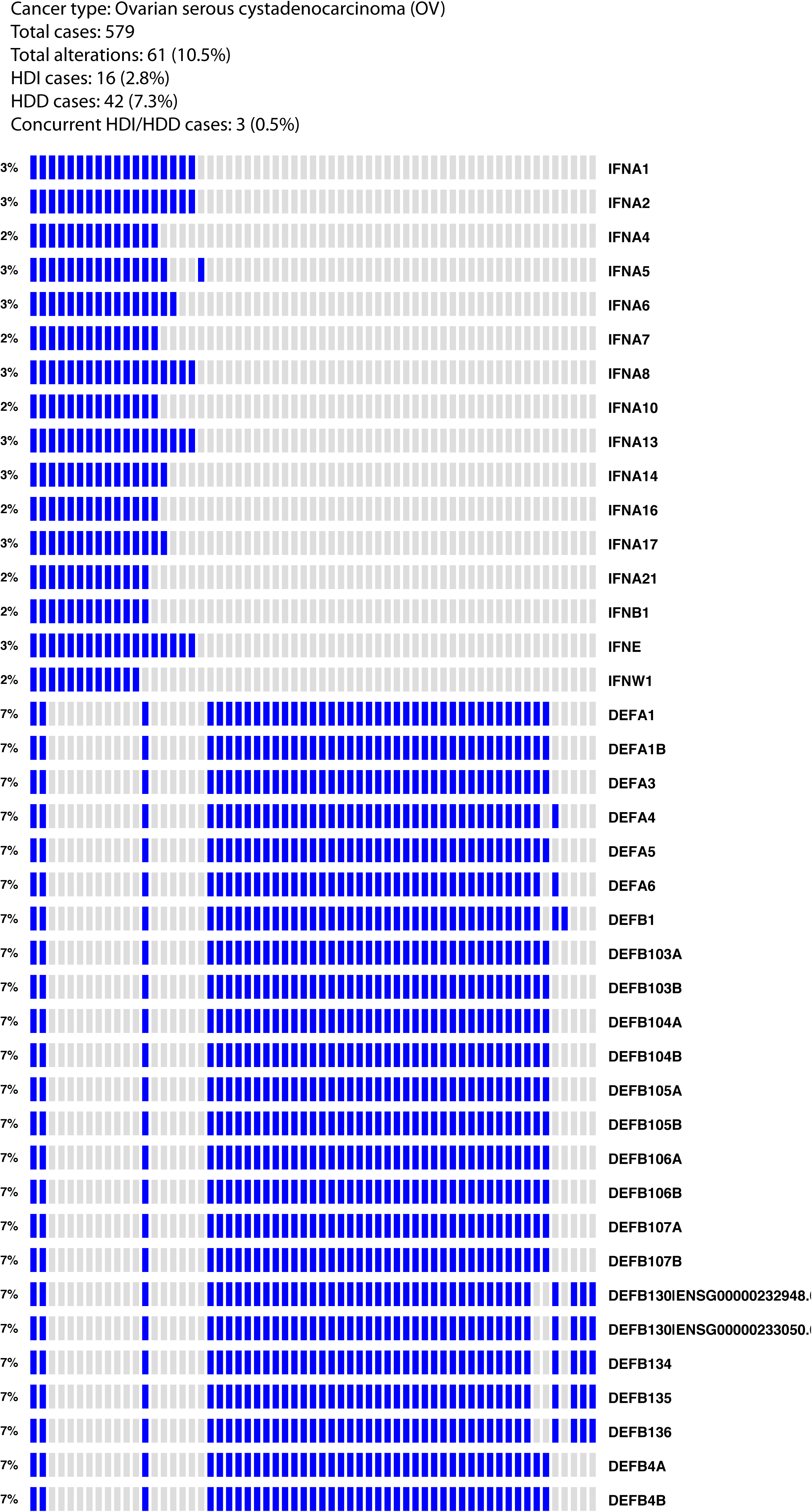
Oncoprint plots for Pancreatic adenocarcinoma (PAAD), Mesothelioma (MESO), Liver hepatocellular carcinoma (LIHC), Uterine carcinosarcoma (UCS), Breast invasive carcinoma (BRCA), Ovarian serous cystadenocarcinoma (OV), Head and Neck squamous cell carcinoma (HNSC), and Skin Cutaneous Melanoma (SKCM).

**Figure S7.**
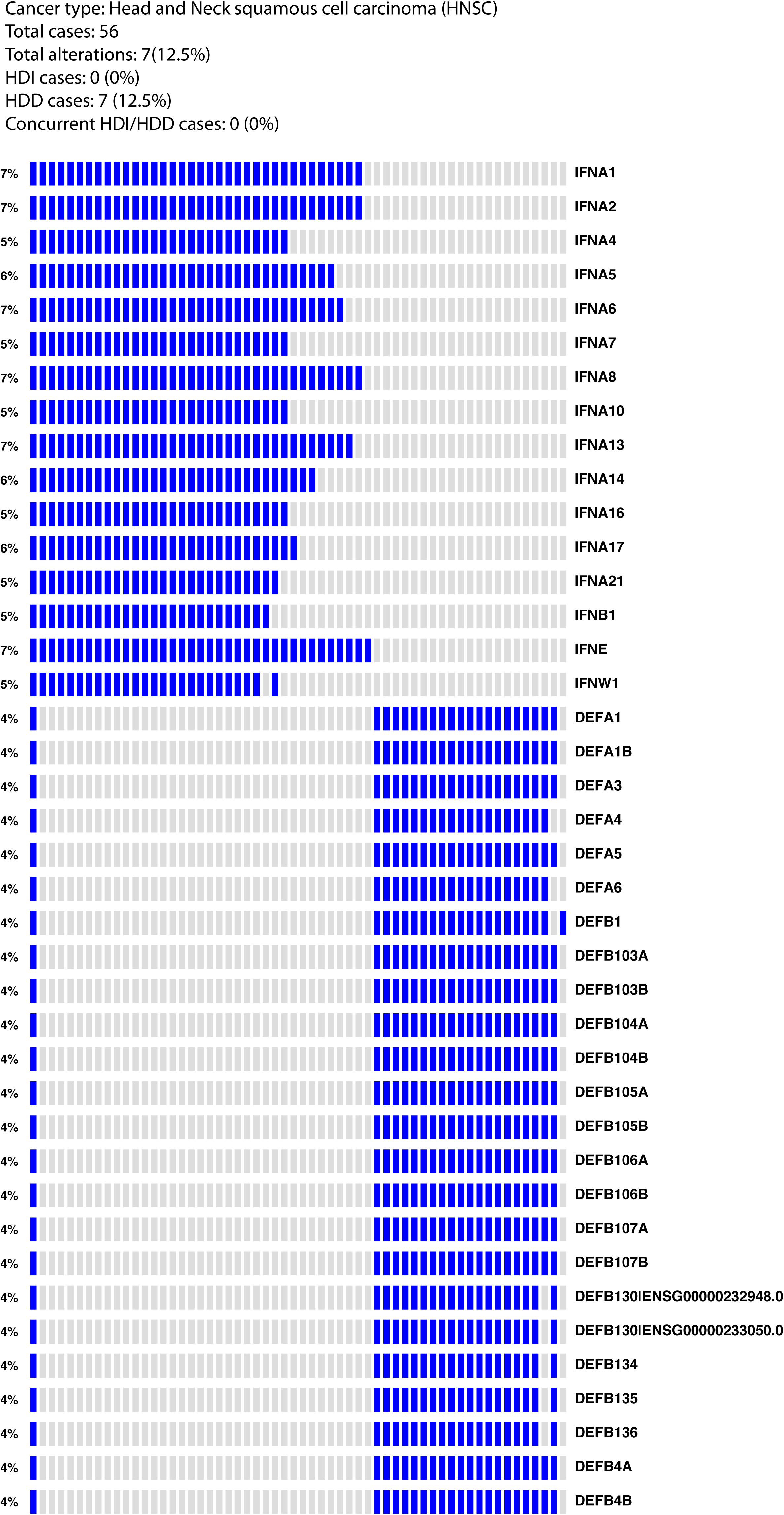
Oncoprint plots for Pancreatic adenocarcinoma (PAAD), Mesothelioma (MESO), Liver hepatocellular carcinoma (LIHC), Uterine carcinosarcoma (UCS), Breast invasive carcinoma (BRCA), Ovarian serous cystadenocarcinoma (OV), Head and Neck squamous cell carcinoma (HNSC), and Skin Cutaneous Melanoma (SKCM).

**Figure S8.**
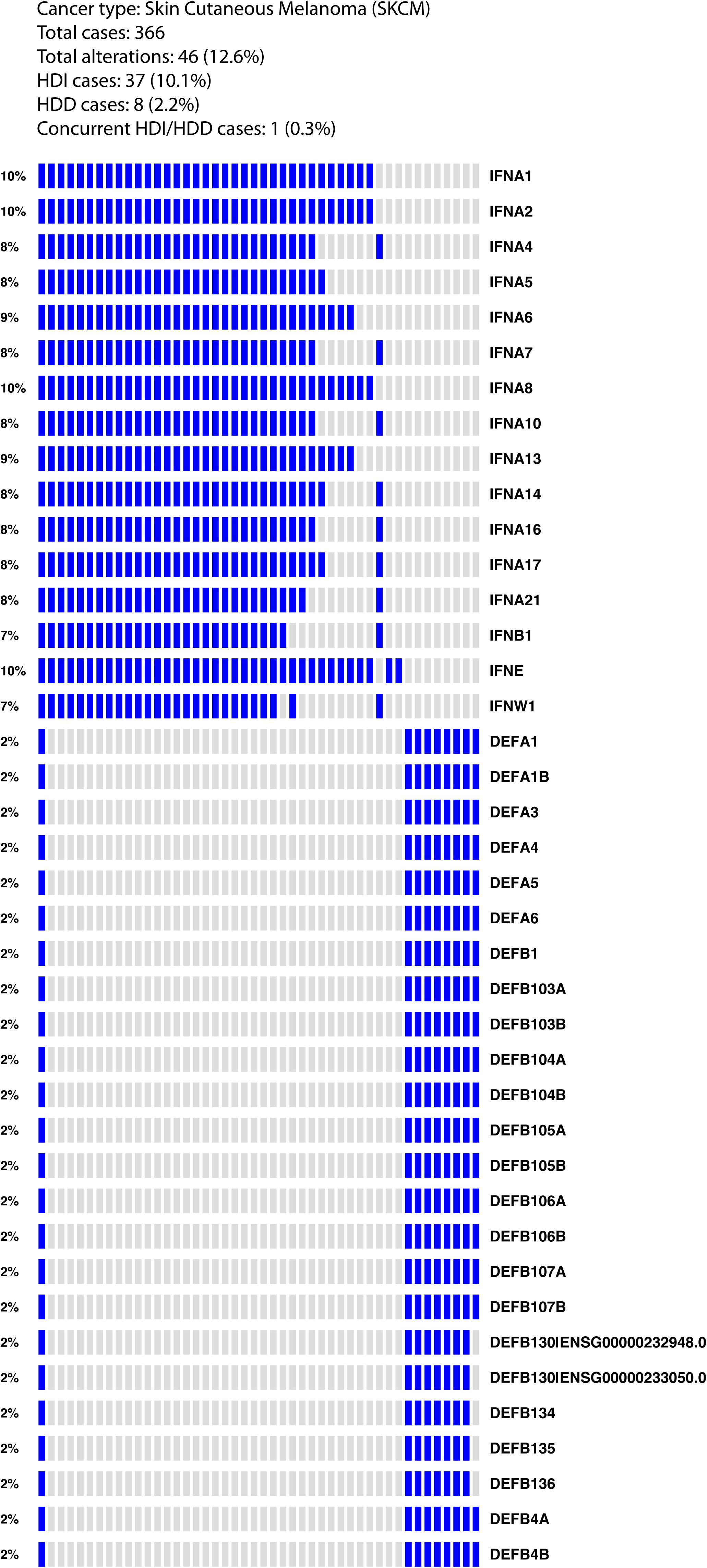
Oncoprint plots for Pancreatic adenocarcinoma (PAAD), Mesothelioma (MESO), Liver hepatocellular carcinoma (LIHC), Uterine carcinosarcoma (UCS), Breast invasive carcinoma (BRCA), Ovarian serous cystadenocarcinoma (OV), Head and Neck squamous cell carcinoma (HNSC), and Skin Cutaneous Melanoma (SKCM).

**Figure S9.**
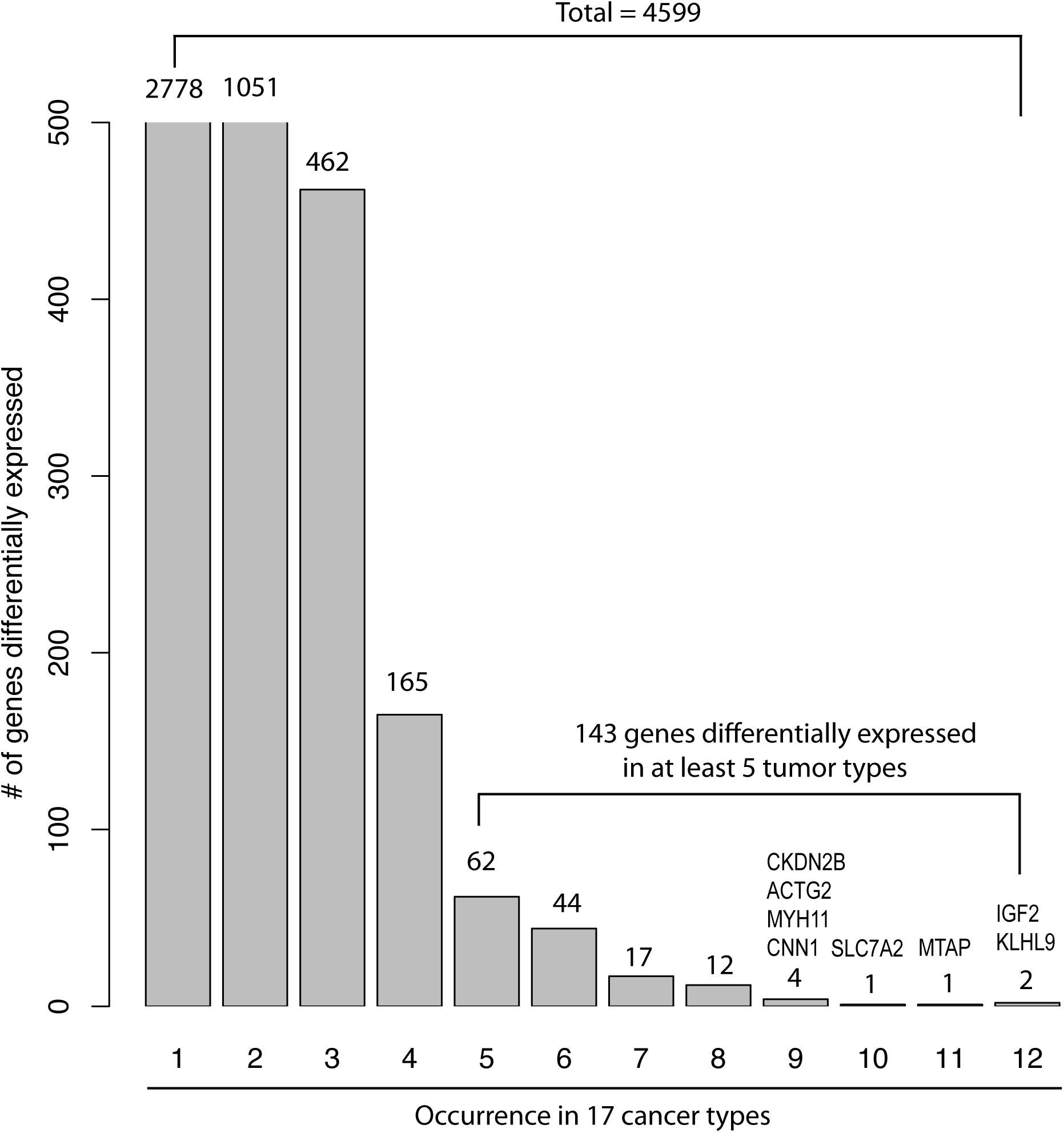
Distribution of 4599 genes that differentially expressed between HDI/HDD positive group and HDI/HDD negative group in at least one tumor type.

**Figure S10.**
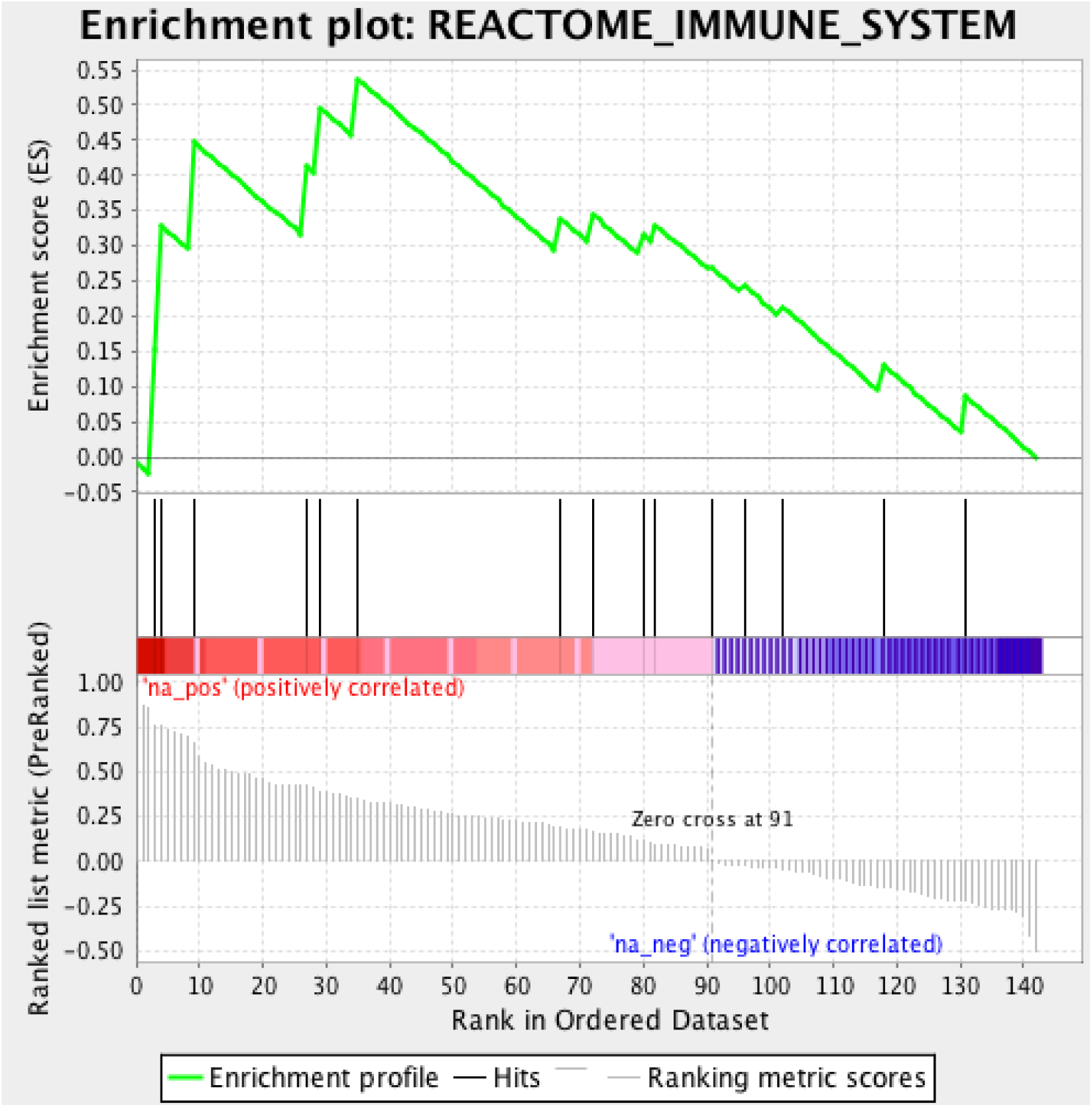
The top enriched gene set detected by GSEA from 143 genes that were commonly altered in at least five tumor types. GSEA, Gene Set Enrichment Analysis.

**Figure S11.**
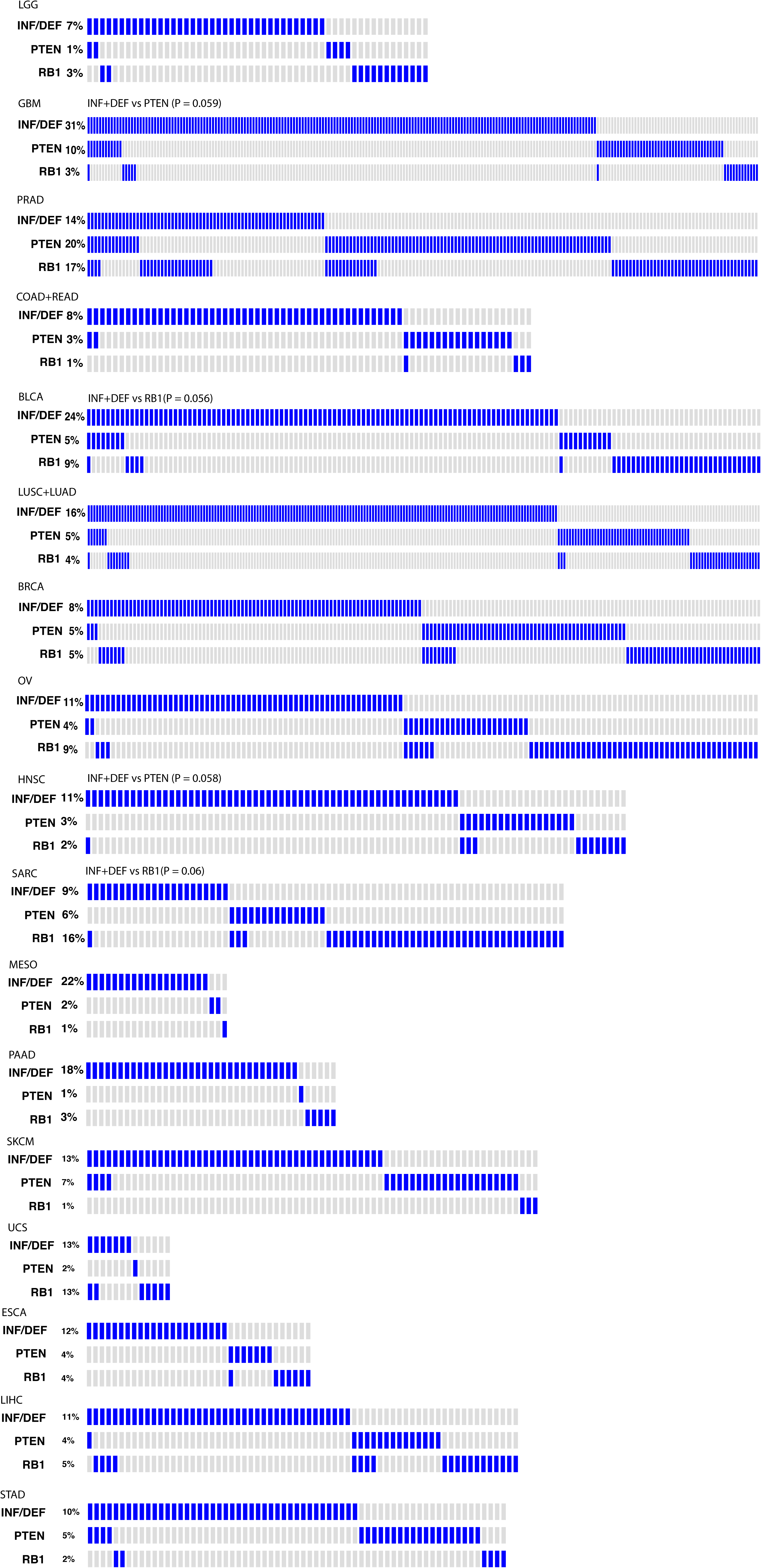
Oncoprint plot showing mutually exclusive patterns between HDI/HDD, PTEN and RB1 homozygous deletions in 17 tumor types.

**Figure S12.**
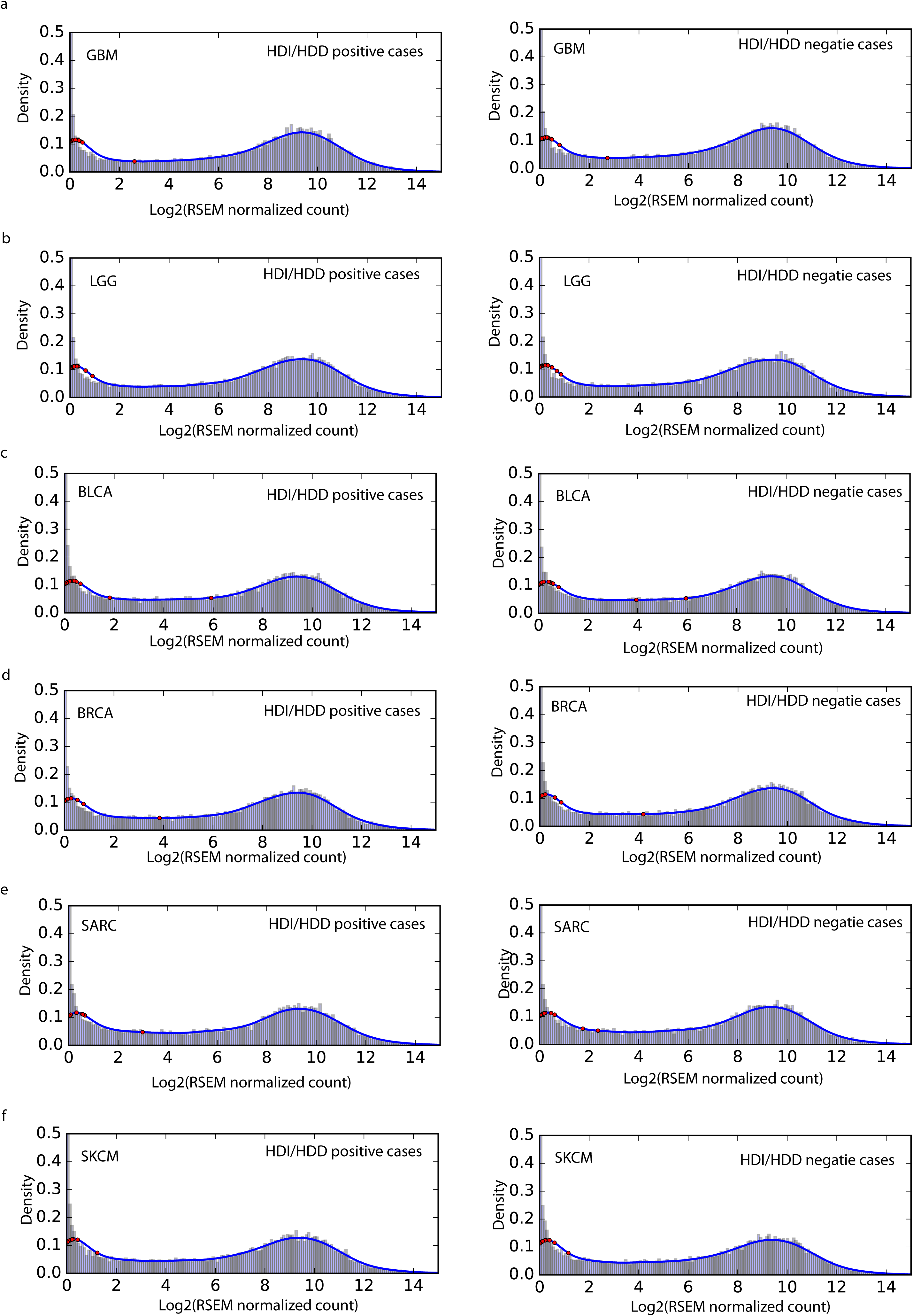
Distribution of gene expression measured by log2 (RSEM normalized read count) for GBM (A), LGG (B), BLCA (C), BRCA (D), SARC (E), and SKCM (F). Red circles indicate interferon and defensin genes.

**Figure S13.**
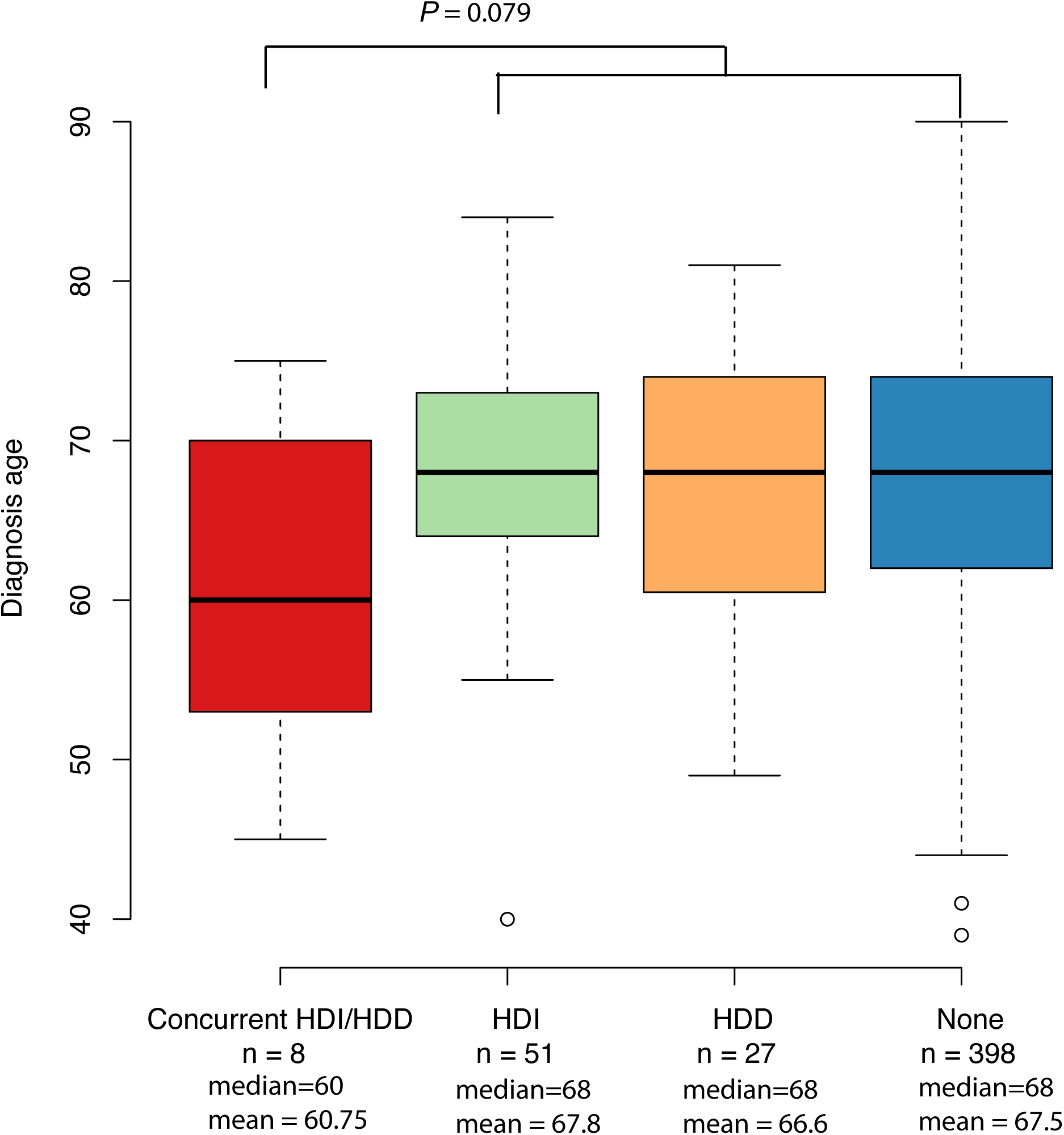
Comparing the median diagnosis ages of LUSC patients who have concurrent HID/HDD (red, n = 8), HDI only (green, n = 51), HDD only (orange, n = 27) and no homozygous deletion (blue, n = 398). The p-value is calculated by Wilcoxon rank sum test (two-sided).

